# Structural and contextual biases interact to shape duration perception

**DOI:** 10.64898/2026.01.07.698096

**Authors:** Laetitia Grabot, Anne Giersch, Pascal Mamassian

## Abstract

Human time perception is flexible and shaped by both structural constraints and contextual influences. Disentangling these sources of bias is essential for understanding the predictive mechanisms underlying temporal perception, yet no unified model currently integrates them. Here, we quantified precisely the structural and contextual biases in a duration discrimination task to constrain models of duration perception. Using a two-interval duration discrimination task, participants judged which of the two stimuli lasted longer. Stimuli were both visual, both auditory, or one of each modality. Consistent with previous findings, auditory stimuli were perceived longer than visual stimuli of equal duration, reflecting intrinsic properties of audiovisual neural processing. Contextual biases were manipulated through different duration distributions from which stimuli were drawn, a procedure known to influence temporal judgments. Our results show that duration discrimination relies on distinct representations of each stimulus distribution and is best explained by a combination of Bayesian inference and rescaling. While Bayesian inference accounts for contextual effects that bring perceived durations closer to the mean of each distribution, rescaling mechanisms have the opposite effects. These findings challenge existing accounts of contextual effects in time perception and suggest that the brain normalizes temporal representations to adapt to environmental statistics.

**Fundings:** This work was supported by the FRC and the UNAFAM association, and scientific expertise is provided by the FRC’s Scientific Advisory Board, and the Anneliese Maier Award to Pascal Mamassian from the Alexander von Humboldt Foundation.

**Data availability:** Behavioral data and modeling codes will be open-access upon publication.

## Introduction

Time does not flow the same way for everyone because individuals weigh sensory modalities and temporal context differently. For example, when watching dubbed movies for an extended duration, some people are sensitive to problems of approximate synchronization between image and sound, while others do not notice anything (Ipser et al., 2017, 2018). These discrepancies in time perception may reflect adaptive mechanisms operating across multiple timescales. Individuals can temporarily adapt to and compensate for physical asynchronies, yet they also exhibit stable, idiosyncratic differences in brain structure, such as the connectivity between timing-related brain areas. Such differences may arise from longer-term adaptive processes. It is important to differentiate two types of prior constraints that shape time perception. Contextual priors refer to information that is valid only in a specific context and that is ignored, or sometimes forgotten, once we move to a different context. In contrast, structural priors correspond to brain constraints that will be effective in all situations (Seriès & Seitz, 2013; Teufel & Fletcher, 2020). Because these two types of priors differ in nature and likely rely on distinct mechanisms, clearly distinguishing between them will be essential for understanding how the brain constructs the experience of time. More generally, this idea has recently been discussed as a general approach (de Lange et al., 2018; Yon et al., 2019; Teufel & Fletcher, 2020) that could bring new insights in understanding the biological implementations of the predictive brain framework. However, while contextual priors have been extensively studied, structural priors remain comparatively underexplored (Seriès & Seitz, 2013; Teufel & Fletcher, 2020). As a first step to fill this gap, we aim here to develop a computational model to quantify the impact of both structural and contextual priors on time perception in humans.

Time perception is surprisingly highly variable and depends on the context and the task in hand, and this variability hinders making progress on major questions about time perception, such as how the brain computes time (Matthews & Meck, 2014). The extensive literature shows that all aspects of time, whether duration, order or mental time travel, are influenced by context-based priors that reflect environmental constraints. For instance, perceived duration strongly depends on the temporal statistics of the environment (Jazayeri & Shadlen, 2010), and perceived simultaneity can expand following an exposure to persistent stimuli asynchrony (Fujisaki et al., 2004). The large lability in time perception has therefore been nuanced (Sanford, 1888; Grabot & van Wassenhove, 2017). Indeed, even if inter-individual variability is large, the characteristics of a given participant remain stable when tested several times, such as for instance their bias to order auditory and visual stimuli (Grabot & van Wassenhove, 2017). This observation led to the idea that perceived simultaneity relies in part on idiosyncratic traits (Grabot & van Wassenhove, 2017; Sanford, 1888). Importantly, these biases are not overcome by attentional manipulation (Grabot & van Wassenhove, 2017) or contextual influences (Grabot & Kayser, 2020), suggesting that structural priors may play an important role in time perception at the individual level. However, time perception is usually studied on the basis of group average data, ignoring individual biases, and thus potentially leading to an overlook of crucial mechanisms.

The present study aims at jointly quantifying the impact of both contextual and structural priors on time perception, by combining psychophysics with individual-level modeling of behavioral data. Participants performed a duration discrimination task, comparing the duration of two stimuli, both visual, both auditory, or one of each (Figure 1). Sounds are usually perceived longer than visual events (Goldstone, 1968; Goldstone & Lhamon, 1972, 1974), which may be due to the intrinsic properties of sensory timing systems (Wearden et al., 1998). Inter-individual differences in this audiovisual bias will be used as a proxy to measure structural priors. The contextual prior was manipulated by changing the distribution of presented stimuli. Temporal estimates are known to depend on the distribution from which they are drawn: the same duration is underestimated if drawn from a pool of short durations, but overestimated if drawn from a pool of long durations (Jazayeri & Shadlen, 2010; Acerbi et al., 2012; Murai & Yotsumoto, 2016; Roach et al., 2017; Maaß et al., 2019; Damsma, Schlichting, & van Rijn, 2021; Damsma, Schlichting, van Rijn, et al., 2021). This reflects a classical central tendency effect, in which perceived durations are biased toward the mean of the overall distribution.

**Figure 1.**
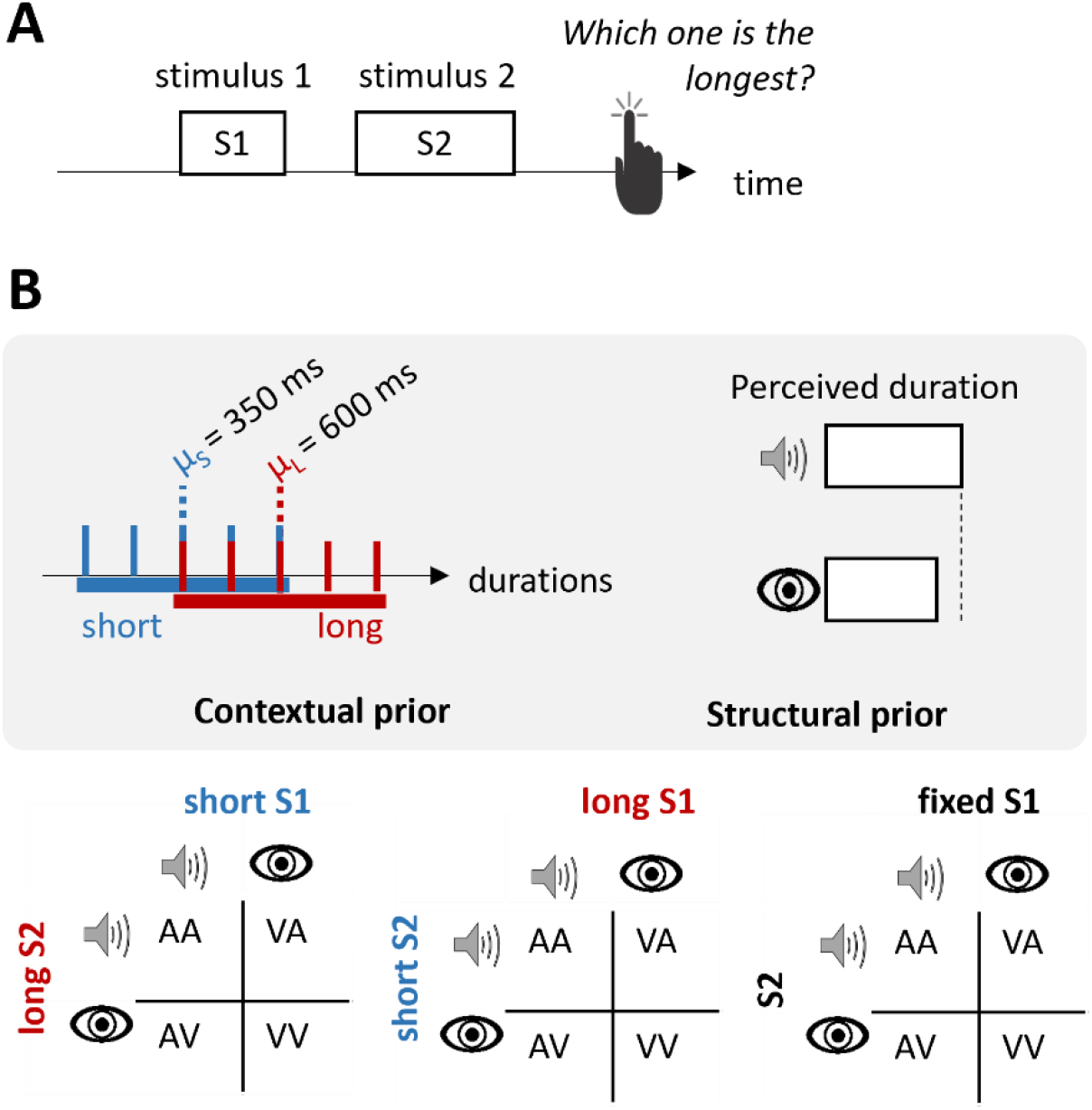
Duration discrimination task involving structural and contextual priors. **A**. Participants judged which of the two presented durations was perceived longer. **B**. The contextual prior was manipulated by altering the distributions from which the first and second stimuli were drawn (short or long distributions, in blue and red respectively, in log space). The structural prior was assessed by comparing participants’ perceptions of the duration of visual and auditory stimuli with identical physical durations. All combinations of conditions were tested, as well as four supplementary conditions where the duration of the first stimulus was fixed.

The durations perceived by each participant, as well as the inter-individual differences in performance will be modeled using tools of Bayesian theory (Figure 2A). According to this probabilistic framework, perception arises from a combination of noisy sensory information and prior knowledge. The main components of Bayesian models are the likelihood function (the probability distribution reflecting the uncertainty in the measured duration given the physical duration), the prior probability distribution (knowledge of the temporal context), and the posterior distribution obtained by combining the first two (estimation of the physical duration given the measurement). Bayesian models have successfully explained the change of performance in a temporal task due to contextual priors (Jazayeri & Shadlen, 2010; Mamassian & Landy, 2010). The main novelty of our approach is to consider two sources of priors in the model (structural and contextual), and test how they are combined, by assessing their respective weight in the final duration estimate.

**Figure 2.**
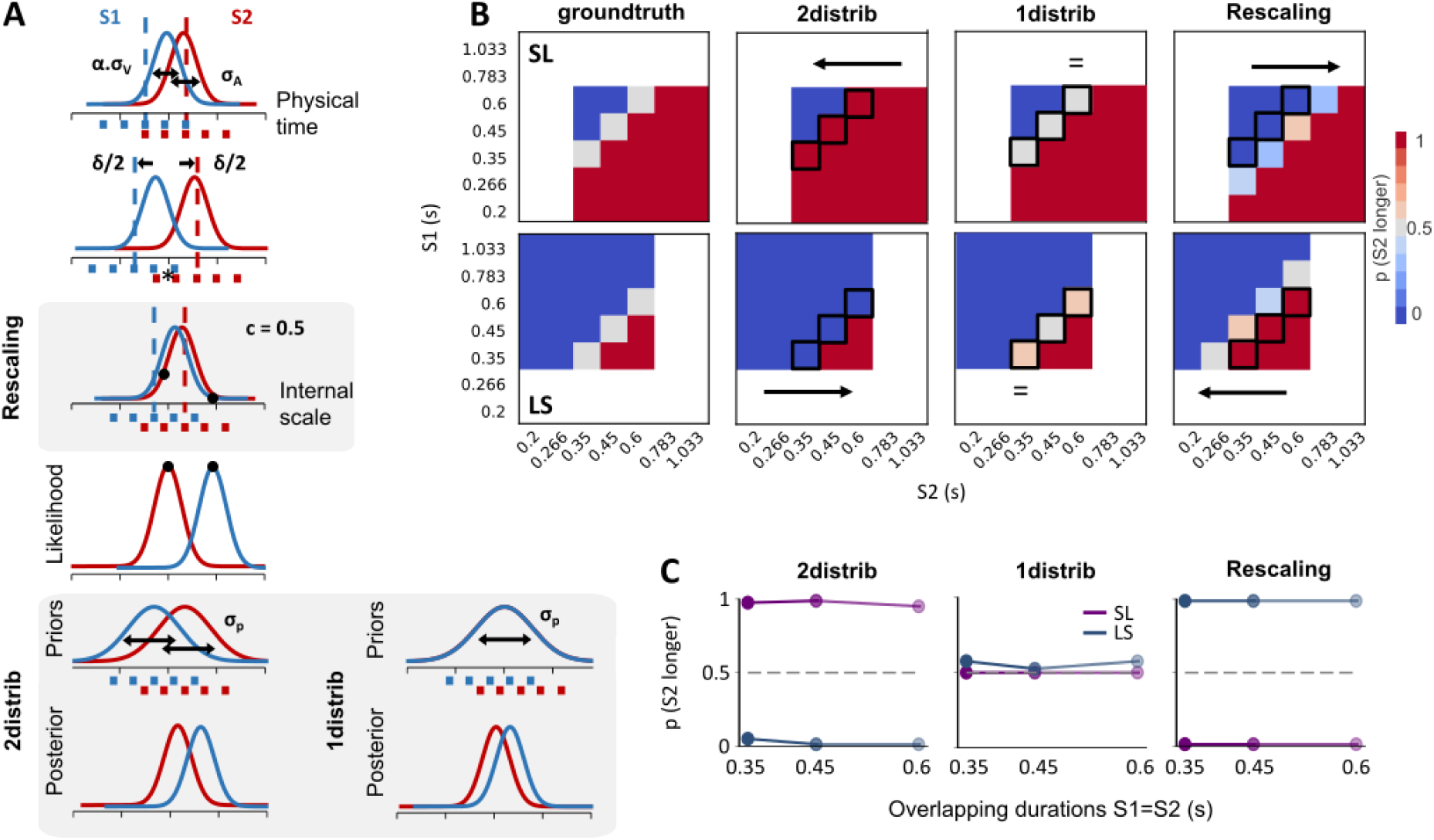
Behavioral models and predictions. **A**. Three distinct implementations of contextual biases were developed as separate models. For each trial, participants have to compare two durations (S1 and S2) presented in two consecutive intervals. They generate some internal representations whose uncertainty is modelled as Gaussians centered on the physical duration. If a visual stimulus is presented in the first interval and an auditory stimulus in the second, the standard deviations of these distributions are (α.σ_v_) and σ_A_, respectively (the parameter α characterizes the fact that the first interval might be more uncertain by the time the perceptual decision is taken). Each distribution is shifted by δ/2 to account for the structural prior (difference of perceived durations between modalities). For models that include “rescaling”, the uncertainty distributions are shifted one more time to realign them towards their barycenter (see position of the asterisk). We illustrate here a partial rescaling (c=0.5); a full rescaling would completely realign the two distributions. The posterior probability is calculated using the likelihood and prior probability. In the “2distrib” models, the distributions from which S1 and S2 are drawn are represented separately, while a wider single distribution is used in the “1distrib” models. **B**. We consider models that differ in the use of the “2distrib” or “1distrib” contextual priors, and the use of “rescaling”. These three components generate distinct patterns of predicted duration discriminations, especially when one compares the blocks short-long (SL, top row) and long-short (LS, bottom row) to the ground truth (left column). In these illustrations, the model parameters took arbitrary values (δ = 0.4, σ_A_ = 0.05, σ_V_ = 0.08, σ_c_ = 0.1, c = 0,.5 α = 1), and the ground truth was calculated based on the comparison of the physical durations. The black arrow or “equal” sign indicates how the pattern of “S2 longer” responses shifts. **C**. The model predictions diverge most strongly for durations that overlap between the S1 and S2 distributions, as illustrated in the three panels which display the data highlighted by black contours in the matrices in Figure 2B.

We considered several implementations of the model to test specific hypotheses on how durations are represented and processed during a discrimination task. Because the two stimuli to be compared were drawn from distinct distributions, our first question was whether the perceptual system maintains separate representations for each distribution or instead encodes a single, broader distribution encompassing all possible durations. The results supported the former: each stimulus distribution appeared to be distinctly represented and this impacted behavior. Next, we tested whether the behavioral data were purely driven by Bayesian processes predicting the central tendency effect. Alternatively, we considered whether an additional recalibration mechanism would better account for the data (Fujisaki et al., 2004), given that Bayesian and recalibration processes yield opposite predictions (Figure 2B and 2C). After repeated exposure to a constant lag, the perceptual system might compensate for this lag, effectively rescaling the temporal representation and shifting the perception of subsequent durations (Miyazaki et al., 2006; Murai & Yotsumoto, 2016). Our results indicated that the best-fitting model required both Bayesian and rescaling processes. Finally, we included in the models a parameter to capture a specificity of our experimental paradigm. Because participants have to compare the duration of two stimuli presented sequentially, we allowed for an ordering effect whereby the first stimulus may be represented more noisily than the second because it must be retained in memory (Ellinghaus et al., 2024a; Raviv et al., 2012).

## Materials & Methods

### Participants

31 participants (19 female, age M = 27 years, SD = 5) took part in the study. All had normal or corrected-to-normal vision, and no known neurological or psychiatric disorder history. All participants were naive as to the purpose of the study, provided written informed consent and were compensated for their participation. The study was performed at the Ecole Normale Supérieure (ENS, Paris, France), in accordance with the Declaration of Helsinki (2013) and approved by the French Ethics Committee on Human Research (N° 2022-A00963-40). One participant only performed one experimental session and was therefore excluded, leaving a total of 30 participants considered for the analyses.

### Stimuli

The experiment was written in MATLAB (Version R2023a; The MathWorks, Inc., Natick, MA) using Psychtoolbox toolbox Version 3.0.19 (Brainard, 1997; Kleiner et al., 2007). The visual stimuli were pink noise annuli (inner diameter = 1°VA, outer diameter = 4°VA, contrast = 0.5) centered on the middle of the screen, presented on a gray background (refresh rate = 120 Hz, resolution = 1920*1080 pixels). The auditory stimuli were pink noise (60 dB, 48 kHz, with 5 ms fade-in and 5 ms fade-out), presented with a sound bar below the screen. They were created with the Binaural toolbox (Akeroyd, 2017). The stability of stimulus timing was verified with an oscilloscope and a photodiode. Auditory and visual 200 ms-duration has a standard deviation of 0.7 and 0.3 ms, respectively (across 10 repetitions), ensuring stable and reliable presented durations.

### Experimental procedure

Participants performed a duration discrimination task in a darkened room, with their head on a chin rest located 85 cm away from the computer screen. During a trial, two stimuli were presented sequentially and participants were asked to judge which one lasted longer (Figure 1A). They replied with a button press; the button mapping was pseudorandomized between participants. The delay between the first stimulus S1 and the second S2 was 0.2 s, and the inter-trial interval lasted between 0.9 and 1.1 s (randomly distributed). Participant were asked to fixate a central fixation cross during the whole experiment.

There were 8 main different conditions varying in both the modality and the distribution from which each stimulus was sampled (Figure 1B). The pair of stimuli were either both auditory, both visual, or one auditory and the other visual (denoted AA, VV, AV and VA). Depending on the condition, the duration of a stimulus in one interval could be drawn from either a short (0.2, 0.267, 0.35, 0.45, 0.6 s) or a long distribution (0.35, 0.45, 0.6, 0.783, 1.033 s). The five items of these distributions were placed on a logarithmic scale. Their median values were 0.35 s and 0.6 s, which corresponds to a ratio of 1.7, shown to be large enough to create two distinct prior representations (Maaß et al., 2019). We tested all possible combinations for S1 coming from one distribution, and S2 coming from the other distribution, hence resulting in 8 “main” conditions (denoted SL-AA, SL-VV, SL-AV, SL-VA, LS-AA, LS-VV, SL-AV and SL-VA conditions). Each S1-S2 pair was repeated 10 times.

To assess the structural prior independently of the contextual prior, we added 4 “baseline” conditions designed. To make the contextual prior irrelevant, in these conditions the first interval had a fixed duration, and the duration of the second interval was drawn from a distribution whose median equated the duration of the first interval. Therefore, the contextual prior was similar for both stimuli and thus canceled out when the stimuli were compared, and participants had to register only one distribution. S1 was fixed to 0.45 s and S2 was sampled from a distribution that was larger than those used in the 8 main conditions, covering both the short and long distributions (7 values ranging from 0.2 to 1.033 s). The four combinations of stimuli and modality were tested (AA, VV, AV, VA), with 10 repetitions per duration.

The experiment was split into two identical sessions to ensure it remained manageable in duration (∼1h20 per session, breaks included). At the beginning of the session, participants were trained to the task during ∼5 min with the four baseline conditions (S1-fixed) using only the most extreme durations (0.2 and 1.03, 3 repetitions per duration). They received feedback after each trial (color of the fixation cross changing to green if correct, red if incorrect). Feedback was then removed for the remaining of the experiment. Participants then performed the 4 blocks of baseline task, presented in pseudorandom order. After a break, they performed 8 blocks of the main task (one condition per block, e.g. SL-AA, SL-VV, SL-AV…, Fig. 1B, lower panel), presented in pseudorandom order.

### Behavioral models

To quantify both contextual and structural priors, we started with Bayesian models to estimate the behavioral response of participants to the duration discrimination task because these models are specifically designed to process prior information. In addition to Bayesian processes, we also considered the possibility of a rescaling of the durations of both intervals onto a common internal dimension. The idea behind this rescaling is that the perceptual system attempts to shift the mean of experienced durations to a single reference value (e.g. 0) so that each stimulus is evaluated relative to that reference. This realignment reflects a recalibration of the perceptual system to the range of relevant durations, similarly to the way the visual system adapts to different ranges of luminance when we leave an indoor environment and move to outdoor in full sun light.

Altogether, we considered five models, each testing different hypotheses underlying the implementation of contextual priors (Figure 2A). These models take into account a prior on the distribution of durations or not (“2distrib” or “1distrib” labels in the name of the model below), and a rescaling process or not (“rescaling” label in the name of the model). The “2distrib” component posits that the perceptual system maintains separate representations of the distributions of the first and second intervals, from which the first and second durations are drawn. In contrast, the “1distrib” component assumes that the system relies on a single unified distribution encompassing all possible durations experienced in this block of trials. Lastly, the “rescaling” component brings an additional step where both distributions are first realigned onto a common internal axis before being used in the Bayesian estimation. We hypothesize that the realignment can be done separately for the first and second intervals. All the models we considered included a structural prior, i.e. the extent to which a participant overestimates a sound compared to a visual stimulus of the same duration.

In the duration discrimination task, participants are presented with two sequential stimuli and have to judge which duration is the longest. The “2distrib” models estimate durations as follows. A duration *t*_*s*_ is represented in log space by a Gaussian distribution centered on the true physical duration *t*. If the stimuli are not from the same sensory modality, the Gaussian distributions are shifted by δ, a free parameter capturing the structural prior. The standard deviation of the Gaussian is a free parameter, σ_*A*_ or σ_*v*_, depending on the modality of the stimulus. Therefore, the distribution for an auditory stimulus is

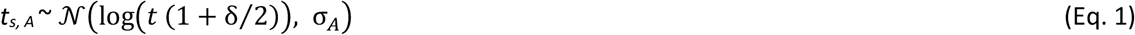

and for a visual stimulus, it is

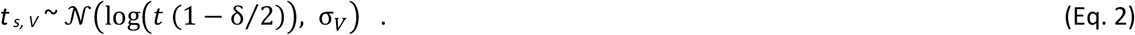

By the time of the perceptual decision, the representation of the stimulus that was presented in the first interval might be noisier than the one in the second interval due to the necessity to maintain the first duration in memory (Raviv et al., 2012; Ellinghaus et al., 2024b). To account for this ordering effect, we introduce an additional parameter α that scales the standard deviation of the first stimulus. Therefore, the equations above correspond to the stimuli presented in the second interval, and to obtain the distributions for stimuli presented in the first interval, the standard deviations are replaced by (*α*.σ_*A*_) or (*α*.σ_*v*_).

Under the assumption that the perceptual system has a noisy representation of a given physical duration, a duration *t*_*m*_ is sampled from the Gaussian distribution defined above to generate the likelihood probability of *t*_*m*_ knowing *t*_*s*_, as a Gaussian centered on *t*_*m*_, with a standard deviation of σ_*A*_or σ_*v*_(or (*α*.σ_*A*_) or (*α*.σ_*v*_) if the stimulus is in the first interval). The prior probability *p* is the knowledge encoded in the perceptual system about the durations distribution. It is represented in log space by a mixture distribution composed of N Gaussian distributions with equal weights (*w*_*n*_ = 1/N), each centered on the log mean of one of the N durations *d*_*n*_ of the stimulus distribution, and with identical standard deviation σ_*c*_ (N = 5 for the main conditions under the “2distrib” component, 7 for the baseline conditions and the main condition under the “1distrib” component). For simplicity, the mixture distribution is approximated by a Gaussian distribution with mean denoted *t*_*p*_ and standard deviation σ_*p*_

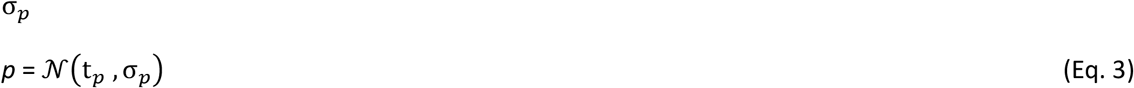

where the mean is

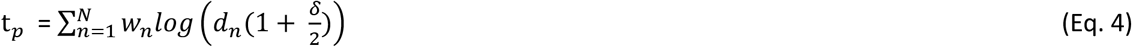

and the standard deviation is computed from the law of total variance

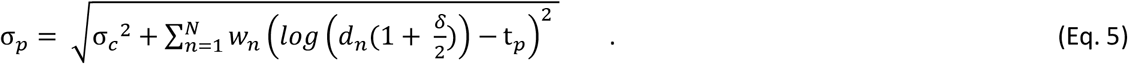

The structural prior also influences the prior probability, which accounts for the presence of the δ term in Equation 4 and 5. Note that these equations are formulated for a distribution of auditory stimuli; for a distribution of visual stimuli, the sign of the δ term is reversed.

The posterior probability is calculated as the product of the likelihood probability with the prior probability, according to Bayes’ rule. This gives a Gaussian with a mean t_*e*_ and a standard deviation σ_*e*_ defined as follows

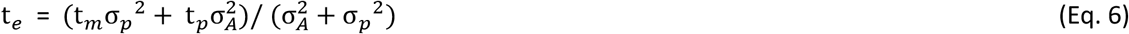

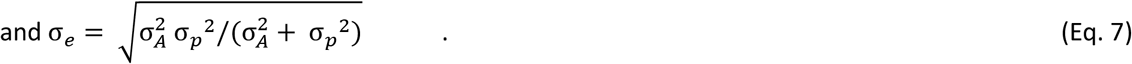

The estimated duration is finally calculated as the duration minimizing the posterior expected loss, using Bayesian least square estimation (Jazayeri & Shadlen, 2010). The output of a given trial is given by estimating the durations of the first and second stimuli and by comparing the two estimates.

All models follow a similar sequence of steps, but with specific variations. The models with a label “2distrib” considered as prior only the distribution from which the stimulus duration to estimate was drawn (i.e. the short or the long distribution). In contrast, the models with a “1distrib” label consider all possible durations (coming from both first and second intervals distributions) in the calculation of the prior. Because the distributions of the first and second intervals overlap partially, the “1distrib” models have weights *w*_*n*_ used in Equation 4 and 5 that are 2/(N +q) for the overlapping durations, and 1/(N+q) for the others (with N = 7). The “rescaling” models have a supplementary first step where the Gaussians representing the prior and the stimulus are shifted toward the barycenter of the combined distributions of the first and second stimuli (composed of M durations, M = 7). The free parameter *c* indicates the extent of rescaling (0: none, 1: full realignment of the two distributions). Equation 1 is therefore rewritten as

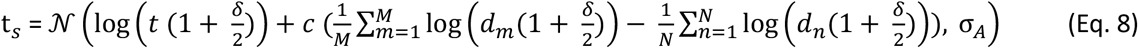

and Equation 4 becomes

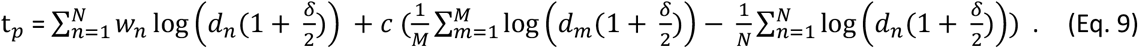

## Models fitting

In total, we tested 5 models that include or not the three components presented above: “2distrib”, “1distrib”, “rescaling+2distrib”, “rescaling”, and “rescalingAV+2distrib”. The “rescaling+2distrib” is a model combining the representation of two distinct distributions for each stimulus to compare, with a first rescaling of these distributions. The “rescaling” model does not use the contextual prior probability to compute the posterior, which is therefore equal to the likelihood. In order to test whether rescaling would be an adaptive mechanism to adjust specifically for cross-modal discrepancies (Fujisaki et al., 2004), the “rescalingAV+2distrib” model only applies the rescaling step to bimodal pairs of stimuli (auditory-visual or visual-auditory), but not to unimodal pairs.

There were six free parameters: the structural prior (δ), the standard deviation of the uncertainty for each sensory modality (σ_*A*_ or σ_*V*_), the standard deviation used to calculate the contextual prior (σ_*c*_), the interval order effect parameter (*α*), and the rescaling parameter (c). We use the negative log likelihood (*NLL*) as the cost function to fit these parameters to the data. The optimization of the models was done using Bayesian adaptive direct search (Acerbi & Ma, 2017) implemented in the PyBADS toolbox (Singh & Acerbi, 2024). This approach alternates between fast and local Bayesian optimization steps and slower exploration of the parameters space, and is particularly adapted when the objective function is stochastic. The maximal number of function evaluations was set to 350, with the final 30 evaluations serving as repetitions of the final solution, used to compute the average parameter values. For each parameter, we specified a maximal and minimal bounds (δ = [-0.9, 0.9], σ_A_ = [0.06, 1], σ_V_ = [0.06, 1], σ_c_ = [0.06, 1], c = [0, 1], α = [0.5, 3]) as well as plausible lower and upper bounds (δ = [0, 0.4], σ_A_ = [0.07, 0.2], σ_V_ = [0.07, 0.2], σ_c_ = [0.07, 0.2], c = [0.05, 0.6], α = [1, 1.5]).

The initial parameters of each model were set after fitting data averaged across all participants, using initial parameters estimated from a grid search. The grid search explored the parameter space by evaluating all possible combinations of four values for each parameter (δ = [0.05, 0.1, 0.2, 0.3], σ_A_ = [0.09, 0.15, 0.2, 0.25], σ_V_ = [0.09, 0.15, 0.2, 0.25], σ_c_ = [0.09, 0.15, 0.2, 0.25], c = [0.1, 0.25, 0.40, 0.55], α = [1, 1.1, 1.2, 1.3]). The combination minimizing the negative log likelihood between model and data was chosen as initial parameters in the fit of average data. The resulting estimated parameters were used as initial parameters in the individual fits (“2distrib”: [δ = 0.21, σ_A_ = 0.35, σ_V_ = 0.52, σ_c_ = 0.97, α = 1.17], “1distrib”: [δ = 0.07, σ_A_ = 0.38, σ_V_ = 0.49, σ_c_ = 0.37, α = 1.52], “rescaling+2distrib”: [δ = 0.11, σ_A_ = 0.34, σ_V_ = 0.44, σ_c_ = 0.54, c = 0.52, α = 1.32], “rescaling”: [δ = 0.18, σ_A_ = 0.33, σ_V_ = 0.42, c = 0.31, α = 0.89], “2distrib + rescalingAV”: [δ = 0.19, σ_A_ = 0.33, σ_V_ = 0.48, σ_c_ = 0.88, c = 0.55, α = 1.02]).

### Models’ comparison

These models were independently fitted to the data and compared using Akaike Information Criterion (AIC), calculated as

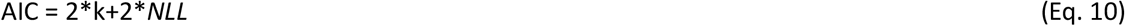

with k the number of free parameters.

### Parameter recovery

To assess the robustness of the model, we performed a parameter recovery analysis that consisted in simulating 1023 datasets from known parameters. The parameters were randomly picked within a 40% range of the final group-average parameters found for the best fitting model (rescaling+2distrib: δ = 0.07, σ_A_ = 0.29, σ_V_ = 0.37, σ_c_ = 0.37, c = 0.65, α = 1.47). Each simulated dataset was then fitted to the “rescaling+2distrib” model (best model found in the model’s comparison analysis) and the recovered parameters were correlated to the known parameters. To investigate potential covariations between parameters, we also correlated between-parameter the deviation of recovered parameter from the known value.

## Results

### Contextual prior: two distributions are represented and rescaled onto an internal axis

We manipulated both contextual and structural priors in a duration discrimination task (Figure 1 and see Figure 3A and 3B for a visualization of the average data). We compared five models based on Bayesian integration, but specifying different implementations of the contextual prior. The “2distrib” component in a model maintains two separate prior durations distributions from which each stimulus is drawn. In contrast, the “1distrib” component in a model imposes that only one single distribution is represented that encompasses all possible durations for S1 and S2. The “rescaling” component in a model provides an additional step in which both distributions are first mapped onto a common internal axis. Figure 2B shows how these different models lead to different predicted behavioral data. Notably, the models’ predictions diverge specifically for the durations that overlap between the S1 and S2 distributions, (that is when the same duration is presented in both intervals). The Bayesian model with a “2distrib” component predicts that the stimulus drawn from the longer-duration distribution will be perceived to be longer (i.e., S1 > S2 in the LS condition and S1 < S2 in the SL condition), consistent with a regression-to-the-mean effect. In contrast, a model with the rescaling component predicts the opposite pattern, as illustrated in Figure 2C. The empirical data show a pattern mostly consistent with the predictions of a rescaling model (Figure 3B, bottom panels). We compared the five models using AIC (Figure 3C). One of the models specifically tested the hypothesis that the perceptual system performs rescaling only when the distributions are cross-modal (“rescalingAV+2distrib”). The best model across participants was the “rescaling+2distrib” model, that is a model combining the representation of two distinct distributions with a rescaling component for all stimuli pairs (ΔAIC summed across participants, for the other four models relative to this winning model were for “2distrib”: 6750, “1distrib”: 1014, “rescaling”: 468, and “rescalingAV+2distrib”: 1487). On an individual basis, this “rescaling+2distrib” model minimized the AIC for 15 participants, the “rescaling” for 9 participants, the “rescalingAV+2distrib” for 4 participants, the “1distrib” for 1 participant and the “2distrib” for 1 participant.

**Figure 3.**
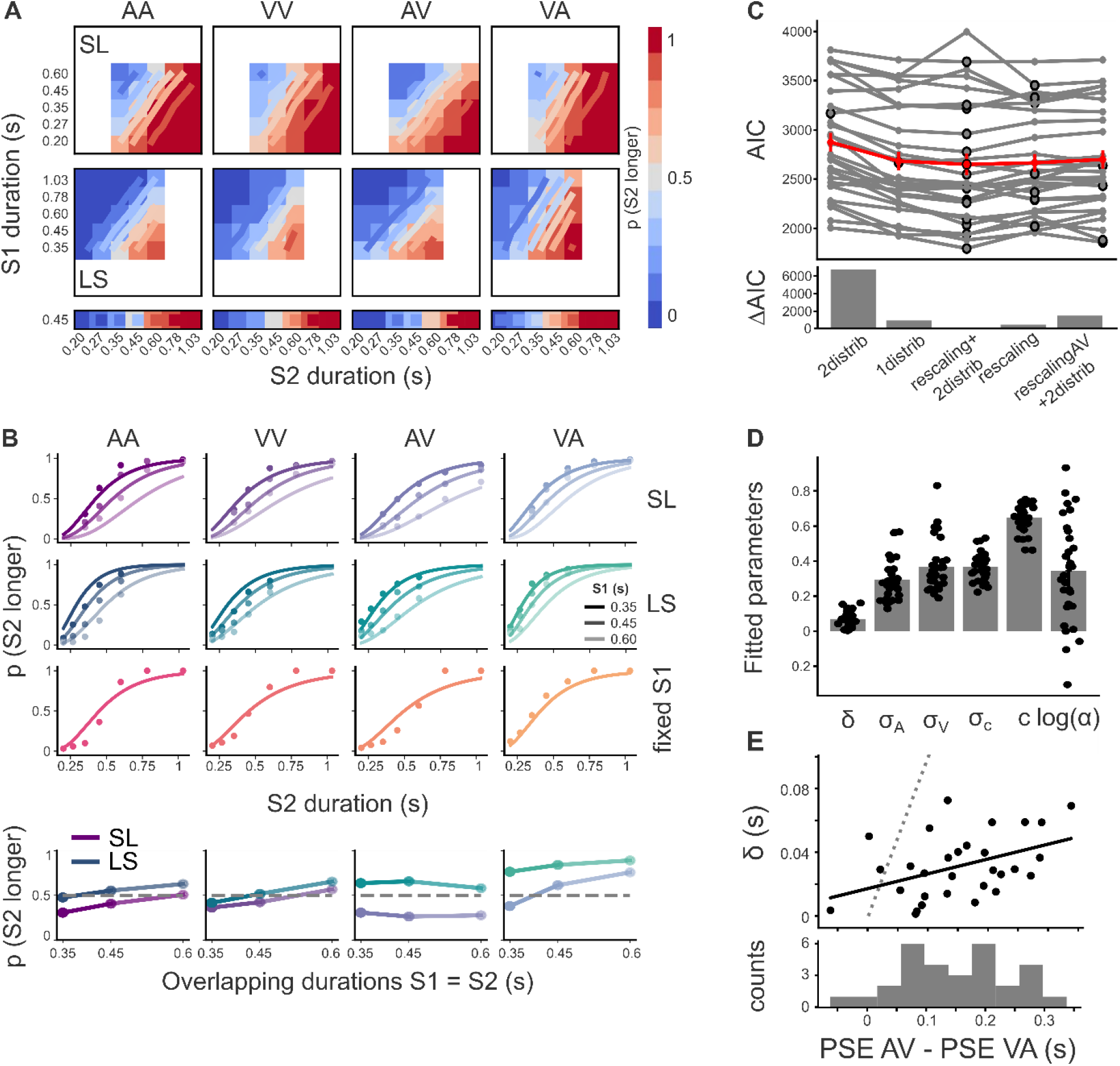
Best model with parameter estimates. The behavioral data are best explained by a model that represents two distinct durations distributions and rescales them onto a common internal axis. **A**. The behavioral data averaged across participants are plotted in matrices using a color scale that represents the probability of perceiving the stimulus in the second interval S2 longer than the one in the first interval S1. Predictions from the best model (“rescaling+2distrib”) are plotted with contour lines superimposed on the matrices with the same color scale. The panels show all pairs of sensory modality comparisons (columns) and each duration distribution across the two intervals (rows, SL= short-long, LS = long-short, and baseline conditions on the bottom row). **B**. The same data are now plotted using psychometric curves. For the SL and LS conditions, only three curves are shown, corresponding to the three S1 durations that were common to the two intervals (0.6, 0.45 and 0.35 s). The dots correspond to the data averaged across participants, while the curves correspond to the best model’s predictions. The four bottom panels reproduce participants’ responses for the overlapping durations (SL in shade of purple, LS in shade of blue), for an easier comparison to the model predictions presented in Figure 2C. **C**. The five models were compared using AIC (Akaike Information Criterion). The best model minimizing AIC is “rescaling+2distrib”. Each line corresponds to one participant, with the minimal AIC highlighted by a large dot. The average is shown in red. In the bottom panel, AICs were first summed across participants and then displayed relative to the “rescaling+2distrib” model. **D**. The fitted parameters of the “rescaling+2distrib” model are shown, each dot representing one participant, and the bar shows the average. **E**. The parameter for the structural prior (δ) correlates significantly with the non-parametric structural prior, defined as the difference between the point of subjective equality (PSE) in the baseline audio-visual condition and the PSE in the baseline visuo-auditory condition (bottom histogram).

The parameters found after fitting the best model to the data are shown in Figure 3D. The uncertainty on the auditory stimuli is significantly smaller than the one on the visual stimuli (mean ± standard deviation: σ_*A*_ = 0.29 ± 0.12, σ_*V*_ = 0.37 ± 0.15, paired t-test: t = -6.06, p = 1.4e-6), which is consistent with previous findings (e.g. Burr et al., 2009; Wearden & Jones, 2021). The parameter representing the contextual prior, that is the standard deviation σ_*c*_, is equal to 0.37 ± 0.08. The rescaling parameter c is equal to 0.65 ± 0.08 indicating a strong but partial rescaling (a full rescaling would have been c = 1, and no rescaling c = 0). Lastly, there is a stimulus order effect, since the parameter *α* is significantly above 1 (*α* = 1.47 ± 0.43, t = 5.92, p =2.0 e-6), indicating that the uncertainty of the stimulus in the first interval is 47% larger than that in the second interval.

### Structural prior: sound duration is overestimated compared to visual duration

The difference of perceived durations between modalities was taken into account in behavioral models as a structural prior. The structural prior, represented by the δ parameter, is on average equal to +0.07 ± 0.04, which corresponds to an average overestimation of the sound of 7%, that is an overestimation of 31 ± 18 ms for an average duration of 450 ms. The parametric structural prior is positive for all participants, showing that they all overestimated the duration of sounds compare to visual stimuli. We correlated this parameter with a non-parametric estimation of the structural prior based on the difference of PSE between the audio-visual and visuo-auditory conditions in the baseline blocks (fixed-S1; Figure 3E, mean ± SD = 152 ± 92 ms, equivalent to about 34%). As expected, we found a significant correlation (R = 0.41, p = 0.023). Note, however, that the correlation did not align with the identity line, with a parametric structural prior lower than the non-parametric one, suggesting that part of the difference in the PSE may have been accounted for by other parameters in our model.

#### Parameter recovery

To test the robustness of the model, we ran a parameter recovery analysis by simulating datasets (N = 1023) with parameters sampled from defined ranges. We fitted the best model (“rescaling+2distrib”) to the simulated datasets and correlated fitted parameters to true known parameters. Each correlation was significant, with a slope close to 1, indicating a good recovery of known parameters (Figure 4A, Table 1). Note that the most strongly biased parameter is σ_*c*_, which is overestimated for small true values of σ_*c*_. The structural prior (δ) is slightly overestimated while the rescaling parameter c is slightly underestimated. To investigate whether these errors might be related, we correlated the deviation from true parameters in between-parameters comparisons (Figure 4B). There is indeed a high negative correlation between δ and c (R = -0.66), which might explain the slight inaccuracies in estimating those parameters. There are also high correlations between the deviations of δ and σ_*c*_ (R = 0.70), σ_*c*_ and c (R = -0.82) and σ_*A*_ and *α* (R = -0.51). All the other correlations were also significant but with a smaller effect size (R<0.4).

**Table 1:** Correlation coefficient, p-value and regression line (slope and intercept) between fitted and true parameters from simulated datasets, for each parameter of the best fitting model.

| | $\delta$ | $\sigma_A$ | $\sigma_V$ | $\sigma_c$ | $c$ | $\alpha$ |
| --- | --- | --- | --- | --- | --- | --- |
| R | 0.41 | 0.87 | 0.84 | 0.43 | 0.86 | 0.88 |
| p | 5.6e-42 | 7.9e-312 | 4.6e-277 | 1.2e-46 | 4.42e-298 | < 10e-100 |
| slope | 0.97 | 0.91 | 0.87 | 0.45 | 0.96 | 0.91 |
| intercept | 0.02 | 0.03 | 0.05 | 0.25 | -0.03 | 0.14 |

**Figure 4.**
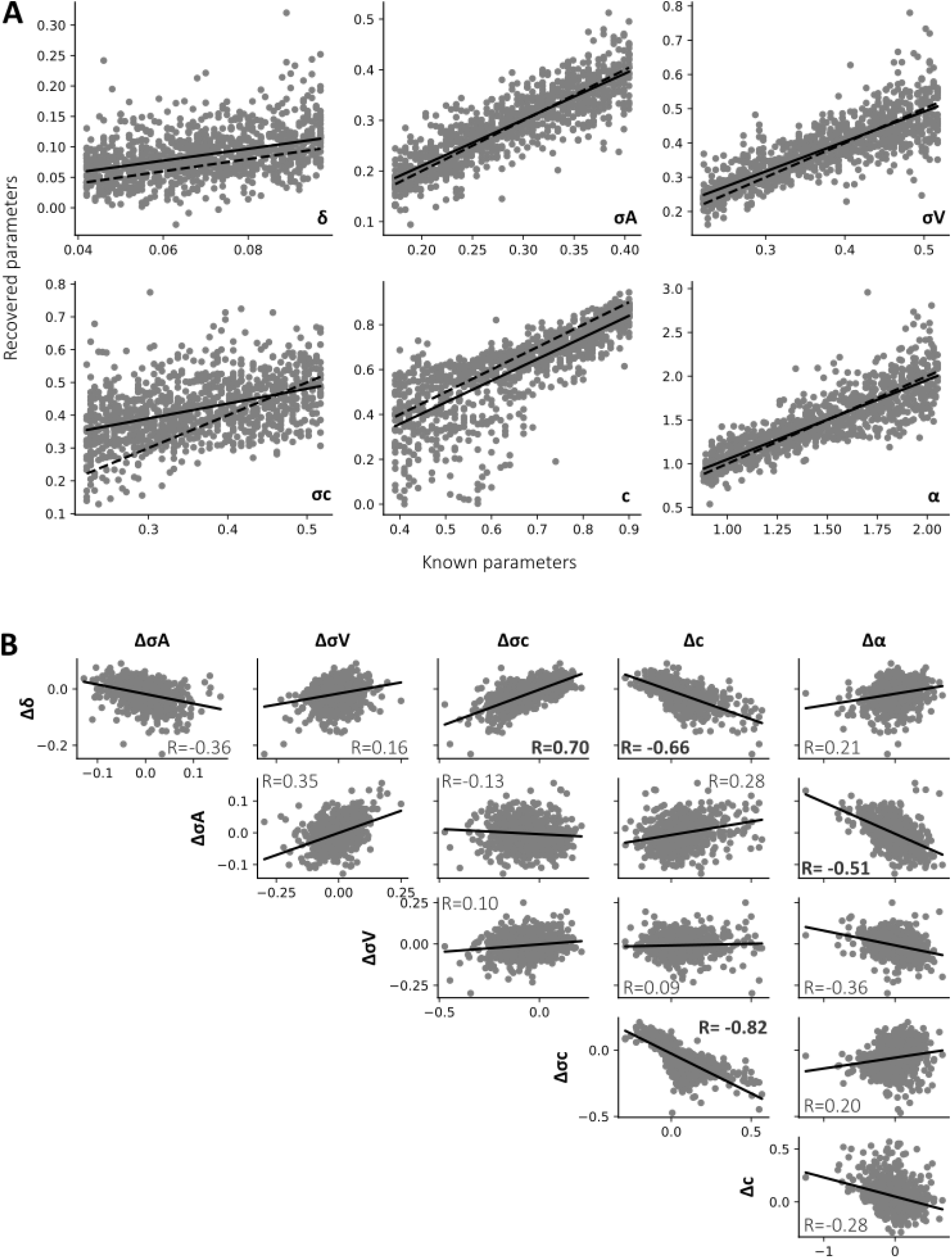
Parameter recovery analysis. **A**. Datasets were simulated by randomly selecting parameters, and fitted to the best fitting model (“rescaling+ 2distrib”) to recover the parameters. The dashed line is the identity line, while the bold line is the correlation between known and recovered parameter. Parameters show good robustness, but with small biases. **B**. To investigate the potential correlation between parameters, we took the differences between simulated and recovered parameters, and correlated these differences between parameters. All correlations (solid line) were significant but the size effects differ: the most significant correlations (R>0.4) are written in bold.

## Discussion

We proposed an experimental paradigm and a unifying model to estimate how both contextual information and structural constraints shape duration perception. When estimating the duration of a stimulus, observers tend to perceive it as closer to the mean duration of the distribution of stimuli that it belongs to rather than its actual duration. This regression-to-the-mean effect is classically well-described by Bayesian models (Jazayeri & Shadlen, 2010; Acerbi et al., 2012; Murai & Yotsumoto, 2016; Roach et al., 2017; Maaß et al., 2019). We adapted these models to a discrimination task, where participants compare two auditory or visual durations presented in succession. By comparing different models, we found that the two distributions from which each presented duration comes from are distinctly represented in the brain. Interestingly, the behavioral data were better explained by a Bayesian model that also included a rescaling component. Rescaling means that before comparing the two durations, the perceptual systems realigned the two representational spaces of each stimulus within a common internal scale, and this scale is used to perform the comparison. This rescaling effect biases the perceived durations in a direction opposite to that of the central tendency phenomenon accounted for by Bayesian theory. Two opposing mechanisms have also been identified in audiovisual temporal order judgments. On one hand, lag adaptation can compensate for a consistent delay between visual and auditory stimuli, while on the other hand, Bayesian integration predicts the opposite effect (Miyazaki et al., 2006; Yamamoto et al., 2012). For both lag adaptation in temporal order judgment and the rescaling process described here in duration discrimination, the perceptual system strives to establish a common perceptual space for integrating diverse sensory inputs.

In our experimental data, the rescaling of both stimulus representational spaces was substantial– around 65%–but not complete. This mechanism accounted for much of the behavioral data, as the Bayesian model alone (the “2distrib” model) provided the poorest fit among those tested. This finding may seem surprising given previous reports of successful Bayesian accounts of duration estimation (Jazayeri & Shadlen, 2010; Acerbi et al., 2012; Murai & Yotsumoto, 2016; Roach et al., 2017; Maaß et al., 2019). We propose that this discrepancy arises from task differences: our study involved a duration comparison task, whereas those previous studies primarily employed temporal production tasks, in which participants reproduced a given interval.

Rescaling can also be understood as comparing each stimulus duration to previously experienced durations within the same temporal interval: the first stimulus is evaluated relative to the prior distribution of first stimuli duration, and the second relative to the prior for second stimuli. It might be a more optimal or less costly strategy to store each duration in a relative code (i.e. as a label “relatively shorter/longer”) rather than in an absolute sensory format (Lee et al., 2013; Teng et al., 2023). The final decision is then based on a comparison of these relative durations.

We initially hypothesized that this strategy would occur primarily when comparing stimuli across modalities; however, this was not supported by our data, as the model applying rescaling only to audiovisual stimulus pairs (“rescalingAV+ 2distrib”) was not the best-fitting model. These results suggest that rescaling may represent a more general strategy employed in discrimination tasks, and further research is needed to test the generality of this mechanism.

Another difference compared to seminal studies on duration estimation (e.g. Jazayeri & Shadlen, 2010) is the order effect induced by the use of two stimuli to compare. Many studies have shown that the discrimination sensitivity is better when the constant standard stimulus precedes the variable one, rather than the opposite (Dyjas & Ulrich, 2014; Bausenhart et al., 2015; Hellström & Rammsayer, 2015; Ellinghaus et al., 2024a). Different hypotheses proposed that this so-called “Type B” effect is due to either the first stimulus being represented as a mixture of a memorized duration combined with recent history, or by a different evidence weighting between the first and second stimuli (Ellinghaus et al., 2024a). In the present study, our experimental design differs from those previous studies because the standard duration was not fixed but varied across trials. Despite equating the variability of the two stimuli, we found that the uncertainty related to the first stimulus was on average about 47% larger than the second stimulus, potentially explained by some memory effect related to the encoding and maintenance of the first stimulus. This effect was observed even though the delay between the two stimuli was short (0.2 s), a duration that has previously been shown to eliminate the Type B effect (Bausenhart et al., 2015; Ellinghaus et al., 2019). This may be explained by the fact that a difference in uncertainty between the two stimuli results in an asymmetrical influence of the prior—stronger for the representation of the first stimulus duration—which leads to a shift in bias but not in sensitivity.

The novelty of our approach lies in examining the impact of structural priors on time perception, in addition to contextual priors. We used the well-established overestimation of auditory relative to visual durations as a good example of structural prior. All participants overestimated auditory compared to visual durations by an average of 7% (approximately 31 ms for a mean duration of 450 ms). This effect aligns with previous reports of auditory overestimation ranging from 6% to 28% (Goldstone, 1968; Goldstone & Lhamon, 1974; Wearden et al., 1998; Penney et al., 2000; Droit-Volet et al., 2004; Ulrich et al., 2006; Droit-Volet et al., 2007; Lustig & Meck, 2011), although it falls in the lower end of that range. Notably, lower estimates in the literature often come from studies using modeling across different conditions (Penney et al., 2000; Droit-Volet et al., 2004; Droit-Volet et al., 2007; Lustig & Meck, 2011), whereas larger effects are typically reported in studies directly comparing behavioral responses (Goldstone, 1968; Goldstone & Lhamon, 1974; Wearden et al., 1998; Ulrich et al., 2006). We further confirmed this effect by correlating parametric and non-parametric estimates of the structural prior, with the latter yielding a higher overestimation (about 34%).

We present a model of temporal discrimination that quantifies both structural and contextual priors, providing a framework for dissecting the underlying neural mechanisms of time perception. Because these biases arise from distinct sources—structural constraints versus contextual adaptation—it is likely that different brain regions and neural oscillatory dynamics contribute to each process (Seriès & Seitz, 2013; Teufel & Fletcher, 2020). This behavioral modeling work represents an essential first step toward uncovering the yet unknown mechanisms by which priors shape our perception.

## Notes

### Competing Interest Statement

The authors have declared no competing interest.

### Summary of Updates

Text clarified, Figure 2 and 3 revised.

## Reference

Acerbi, L., & Ma, W. J. (2017). Practical Bayesian Optimization for Model Fitting with Bayesian Adaptive Direct Search. Advances in Neural Information Processing Systems, 30. https://papers.nips.cc/paper_files/paper/2017/hash/df0aab058ce179e4f7ab135ed4e641a9-Abstract.html

Acerbi, L., Wolpert, D. M., & Vijayakumar, S. (2012). Internal Representations of Temporal Statistics and Feedback Calibrate Motor-Sensory Interval Timing. PLOS Computational Biology, 8(11), e1002771. 10.1371/journal.pcbi.1002771

Akeroyd, M. (2017). A binaural cross-correlogram toolbox for MATLAB [Jeu de données]. The University of Nottingham. 10.17639/nott.320

Bausenhart, K. M., Dyjas, O., & Ulrich, R. (2015). Effects of stimulus order on discrimination sensitivity for short and long durations. Attention, Perception & Psychophysics, 77(4), 1033–1043. 10.3758/s13414-015-0875-8

Brainard, D. H. (1997). The Psychophysics Toolbox. Spatial Vision, 10(4), 433–436.

Burr, D., Banks, M. S., & Morrone, M. C. (2009). Auditory dominance over vision in the perception of interval duration. Experimental Brain Research, 198(1), 49–57.10.1007/s00221-009-1933-z

Damsma, A., Schlichting, N., & van Rijn, H. (2021). Temporal Context Actively Shapes EEG Signatures of Time Perception. J. Neurosci., 41(20), 4514. 10.1523/JNEUROSCI.0628-20.2021

Damsma, A., Schlichting, N., van Rijn, H., & Roseboom, W. (2021). Estimating Time : Comparing the Accuracy of Estimation Methods for Interval Timing. Collabra: Psychology, 7(1), 21422. 10.1525/collabra.21422

de Lange, F., Heilbron, M., & Kok, P. (2018). How do expectations shape perception? Trends Cogn Sci.

Droit-Volet, S., Meck, W. H., & Penney, T. B. (2007). Sensory modality and time perception in children and adults. Behavioural Processes, 74(2), 244–250.10.1016/j.beproc.2006.09.012

Droit-Volet, S., Tourret,Stéphanie, & and Wearden, J. (2004). Perception of the duration of auditory and visual stimuli in children and adults. The Quarterly Journal of Experimental Psychology Section A, 57(5), 797–818. 10.1080/02724980343000495

Dyjas, O., & Ulrich, R. (2014). Effects of Stimulus Order on Discrimination Processes in Comparative and Equality Judgements : Data and Models. Quarterly Journal of Experimental Psychology, 67(6), 1121–1150. 10.1080/17470218.2013.847968

Ellinghaus, R., Bausenhart, K. M., Koc, D., Ulrich, R., & Liepelt, R. (2024a). Order effects in stimulus discrimination challenge established models of comparative judgement : A meta-analytic review of the Type B effect. Psychonomic Bulletin & Review, 31(5), 2275–2284.10.3758/s13423-024-02479-3

Ellinghaus, R., Bausenhart, K. M., Koc, D., Ulrich, R., & Liepelt, R. (2024b). Order effects in stimulus discrimination challenge established models of comparative judgement : A meta-analytic review of the Type B effect. Psychonomic Bulletin & Review, 31(5), 2275–2284. 10.3758/s13423-024-02479-3

Ellinghaus, R., Gick, M., Ulrich, R., & Bausenhart, K. M. (2019). Decay of internal reference information in duration discrimination : Intertrial interval modulates the Type B effect. Quarterly Journal of Experimental Psychology, 72(6), 1578–1586. 10.1177/1747021818808187

Fujisaki, W., Shimojo, S., Kashino, M., & Nishida, S. (2004). Recalibration of audiovisual simultaneity.Nat Neurosci., 7(7), 773–778.

Goldstone, S. (1968). Production and reproduction of duration : Intersensory comparisons. Percept Mot Skills, 26(3), 755–760.

Goldstone, S., & Lhamon, W. T. (1972). Auditory-visual differences in human temporal judgment. Perceptual and Motor Skills, 34(2), 623–633. 10.2466/pms.1972.34.2.623

Goldstone, S., & Lhamon, W. T. (1974). Studies of Auditory-Visual Differences in Human Time Judgment : 1. Sounds are Judged Longer than Lights. Perceptual and Motor Skills, 39(1), 63–82. 10.2466/pms.1974.39.1.63

Grabot, L., & Kayser, C. (2020). Alpha Activity Reflects the Magnitude of an Individual Bias in Human Perception. The Journal of Neuroscience: The Official Journal of the Society for Neuroscience, 40(17), 3443–3454. 10.1523/JNEUROSCI.2359-19.2020

Grabot, L., & van Wassenhove, V. (2017). Time order as psychological bias. Psychological Science, 28(5), 670–678.

Hellström, Å., & Rammsayer, T. H. (2015). Time-order errors and standard-position effects in duration discrimination : An experimental study and an analysis by the sensation-weighting model. Attention, Perception, & Psychophysics, 77(7), 2409–2423. 10.3758/s13414-015-0946-x

Ipser, A., Agolli, V., Bajraktari, A., Al-Alawi, F., Djaafara, N., & Freeman, E. D. (2017). Sight and sound persistently out of synch : Stable individual differences in audiovisual synchronisation revealed by implicit measures of lip-voice integration. Scientific Reports, 7(46413).

Ipser, A., Karlinski, M., & Freeman, E. D. (2018). Correlation of individual differences in audiovisual asynchrony across stimuli and tasks : New constraints on Temporal Renormalization theory. Journal of Experimental Psychology: Human Perception and Performance.

Jazayeri, M., & Shadlen, M. N. (2010). Temporal context calibrates interval timing. Nature Neuroscience, 13(8), 1020–1026.

Kleiner, M., Brainard, D., Pelli, D., Ingling, A., Murray, R., & Broussard, C. (2007). What’s new in psychtoolbox-3. Perception, 36(14), 1–16.

Lee, S.-H., Kravitz, D. J., & Baker, C. I. (2013). Goal-dependent dissociation of visual and prefrontal cortices during working memory. Nature Neuroscience, 16(8), 997–999. 10.1038/nn.3452

Lustig, C., & Meck, W. H. (2011). Modality differences in timing and temporal memory throughout the lifespan. Brain and Cognition, 77, 298–303.

Maaß, S. C., Schlichting, N., & van Rijn, H. (2019). Eliciting contextual temporal calibration : The effect of bottom-up and top-down information in reproduction tasks. Acta Psychologica.

Mamassian, P., & Landy, M. S. (2010). It’s that time again. Nature Neuroscience, 13(8), 914–916. 10.1038/nn0810-914

Matthews, W. J., & Meck, W. H. (2014). Time perception : The bad news and the good. WIREs Cog Sci., 5(4), 429–446.

Miyazaki, M., Yamamoto, S., Uchida, S., & Kitazawa, S. (2006). Bayesian calibration of simultaneity in tactile temporal order judgment. Nature Neuroscience, 9(7), 875–877.

Murai, Y., & Yotsumoto, Y. (2016). Timescale- and Sensory Modality-Dependency of the Central Tendency of Time Perception. PloS One, 11(7), e0158921.

Penney, T. B., Gibbon, J., & Meck, W. H. (2000). Differential effects of auditory and visual signals on clock speed and temporal memory. Journal of Experimental Psychology. Human Perception and Performance, 26(6), 1770–1787.

Raviv, O., Ahissar, M., & Loewenstein, Y. (2012). How Recent History Affects Perception : The Normative Approach and Its Heuristic Approximation. PLoS Computational Biology, 8(10), e1002731. 10.1371/journal.pcbi.1002731

Roach, N. W., McGraw, P. V., Whitaker, D. J., & Heron, J. (2017). Generalization of prior information for rapid Bayesian time estimation. Proceedings of the National Academy of Science, 114(2), 412–417. 10.1073/pnas.1610706114

Sanford, E. C. (1888). Personal Equation. A J Psychol, 2(1), 3–38.

Seriès, P., & Seitz, A. (2013). Learning what to expect (in visual perception). Frontiers in Human Neuroscience, 7. https://www.frontiersin.org/articles/10.3389/fnhum.2013.00668

Singh, G. S., & Acerbi, L. (2024). PyBADS : Fast and robust black-box optimization in Python. Journal of Open Source Software, 9(94), 5694. 10.21105/joss.05694

Teng, C., Kaplan, S. M., Shomstein, S., & Kravitz, D. J. (2023). Assessing the interaction between working memory and perception through time. Attention, Perception & Psychophysics, 85(7), 2196–2209. 10.3758/s13414-023-02785-3

Teufel, C., & Fletcher, P. C. (2020). Forms of prediction in the nervous system. Nature Reviews Neuroscience, 21(4), 231–242. 10.1038/s41583-020-0275-5

Ulrich, R., Nitschke, J., & Rammsayer, T. (2006). Crossmodal temporal discrimination : Assessing the predictions of a general pacemaker-counter model. Perception & Psychophysics, 68(7), 1140–1152.

Wearden, J. H., Edwards, H., Fakhri, M., & Percival, A. (1998). Why « sounds are judged longer than lights » : Application of a model of the internal clock in humans. The Quarterly Journal of Experimental Psychology B: Comparative and Physiological Psychology, 51B(2), 97–120.

Wearden, J. H., & Jones, L. A. (2021). Judgements of the Duration of Auditory and Visual Stimuli. Timing & Time Perception, 9(2), 199–224. 10.1163/22134468-bja10008

Yamamoto, S., Miyazaki, M., Iwano, T., & Kitazawa, S. (2012). Bayesian Calibration of Simultaneity in Audiovisual Temporal Order Judgments. PLOS One, 7(7)(e40379).

Yon, D., de Lange, F. P., & Press, C. (2019). The Predictive Brain as a Stubborn Scientist. Trends in Cognitive Sciences, 23(1), 6–8.

